# Correlation of myeloid-derived suppressor cell expansion with upregulated transposable elements in severe COVID-19 unveiled in single-cell RNA sequencing reanalysis

**DOI:** 10.1101/2023.09.04.556192

**Authors:** Mitra Farahmandnejad, Pouria Mosaddeghi, Mohammadreza Dorvash, Amirhossein Sakhteman, Pouya Faridi, Manica Negahdaripour

**Author notes:** Corresponding authors: Manica Negahdaripour, Pouya Faridi.

## Abstract

Some studies investigated the potential role of transposable elements (TEs) in COVID-19 pathogenesis and complications. However, to the best of our knowledge, there is no study to examine the possible association of TEs expression in cell functions and its potential role in COVID-19 immune response at the single-cell level.

In this study, we reanalyzed single-cell RNA seq data of bronchoalveolar lavage (BAL) samples obtained from six severe COVID-19 patients and three healthy donors to assess the probable correlation of TE expression with the immune responses induced by the SARS-CoV-2 virus in COVID-19 patients.

Our findings indicated that the expansion of myeloid-derived suppressor cells (MDSCs) may be a characteristic feature of COVID-19. Additionally, a significant increase in TEs expression in MDSCs was observed. This upregulation of TEs in COVID-19 may be linked to the adaptability of these cells in response to their microenvironments. Furthermore, it appears that the identification of overexpressed TEs by Pattern recognition receptors (PRRs) in MDSCs may enhance the suppressive capacity of these cells. Thus, this study emphasizes the crucial role of TEs in the functionality of MDSCs during COVID-19.

## Introduction

The rise of severe acute respiratory syndrome coronavirus-2 (SARS-CoV-2) in late 2019 led to a massive crisis and a global pandemic. Since its inception, it has resulted in over 689 million reported cases and more than 6.8 million deaths as of May 31st, 2023 (https://www.worldometers.info/coronavirus/). COVID-19 exhibits a vast spectrum of clinical manifestations based on disease severity in different cases of infection, ranging from mild upper respiratory tract symptoms to severe, multi-organ failure and even death. Although several studies have been done to unravel the detailed mechanistic understanding of COVID-19, the underlying pathogenesis of COVID-19 and its optimal treatment are still far from being clearly understood. (1-3).

Transposable elements (TEs) make up around 50 percent of the human genome and are classified into retrotransposons (class I) and DNA transposons (class II) (4). Retrotransposons, which amplify themselves in the genome, are classified into long terminal repeat (LTR) and non-LTR elements. LTR elements consist of around eight percent of the human genome and are categorized into four families: endogenous retrovirus (ERV), ERV-K, ERV-L, and MaRL. In contrast, non-LTR retrotransposons consist of long-interspersed elements (LINEs), short-interspersed elements (SINEs), and Sine-VNTR-Alu (SVA) (5). It has been debated that dysregulation of TE expression participates in several diseases, from cancer to neurological disorders (5-10).

Recently, some studies have proposed the possible associations between COVID-19 severity and TEs (11-16). Since TEs are activated and involved in inflammatory diseases, various studies suggest the potential role of TEs in COVID-19 pathogenesis. In this regard, Kitsou et al. revealed that TEs are strongly dysregulated in the bronchoalveolar lavage fluid (BALF) samples of COVID-19 patients (12). Besides, Zhang et al. suggested that derepression of LINE-expression, induced by SARS-CoV-2 infection, and consequent inflammatory response, could cause SARS-CoV2 integration into the genome of infected cells (15).

Some studies have been done to decipher the COVID-19 pathology at the single-cell level (17-19) However, there is no single-cell RNA seq study to examine the possible association of TEs dysregulation in cell function and its potential role in COVID-19 immune dysregulation, to the best of our knowledge.

Herein, we reanalyzed RNA-Seq data from the bronchoalveolar lavage (BAL) samples obtained from eight severe COVID-19 patients and four healthy donors at the single-cell level to investigate the potential relationship between TE expression and the immune responses triggered by the SARS-CoV-2 virus in COVID-19 patients.

## Methods

The sc-RNA sequencing data of BAL samples from eight COVID-19 patients (including both alive and dead patients) and four healthy donors were retrieved from the gene expression omnibus GEO (20, 21) database under accession numbers GSE157344 (17) and GSE151928 (22), respectively, and were extensively analyzed. In order to minimize the influence of potential comorbidities, we selected four individuals from each study group who were the most youthful in age. The overall workflow of this method generated by https://biorender.com is depicted in Fig. 1. Moreover, the code is available at git@github.com:mitra-frn/sc-TE-RNA.git.

**Fig. 1.**
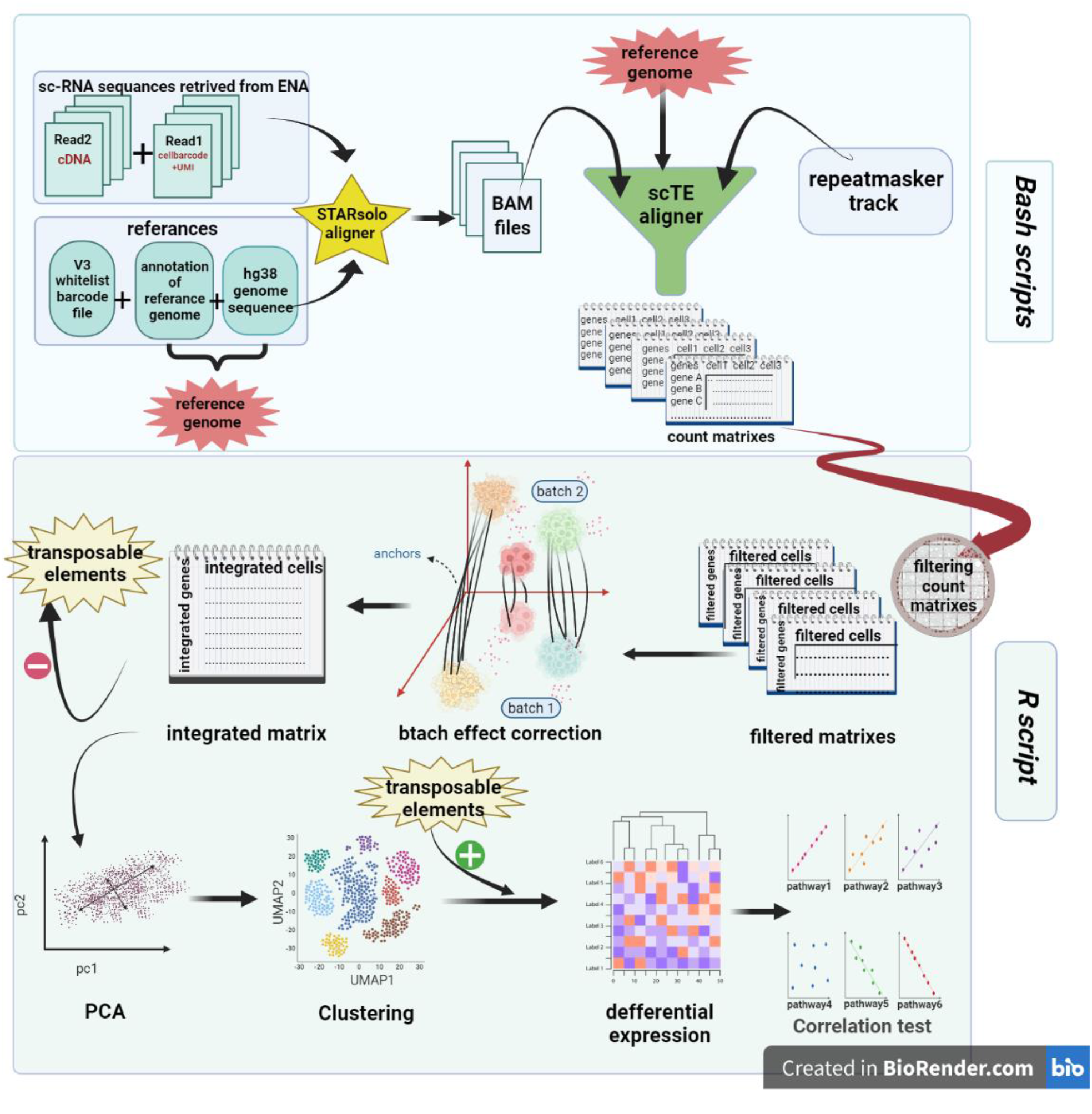
The workflow of this study.

### Aligning reads

Patients’ paired-end FASTQ files were aligned to UCSC hg38 genome assembly using STARsolo (23) and v*3* (3M-february-2018.txt) cell *barcode whitelist files* with the setting’ –outSAMtype BAM SortedByCoordinate,,--winAnchorMultimapNmax 100, --outFilterMultimapNmax 100, --outMultimapperOrder Random, --outSAMmultNmax 1.

The aligned sc-RNA sequence reads of healthy donors’ samples were directly obtained from the ENA database. They were aligned to hg38 genome assembly using STAR aligner as well.

### Quantifying TE expression

The traditional aligners cannot get an accurate quantitation of reads aligned to TEs, as they ignore multi-mapping reads, which is critical for counting TEs. Therefore, we used scTE (24) aligner, which is compatible with STARsolo (23) output files.

Genome indices were built with the *scTE-build* function using UCSC genome browser Repeatmasker track (25) and GENCODE in two modes: exclusive and nointron, and BAM files were realigned via *scTE* function. Then, count matrices made by scTE were applied for further analysis.

### Preprocessing, analysis, and exploration of scRNA-seq data

The Seurat (26-29) is an R package (satijalab.org/Seurat) designed to analyze count matrixes and visualize data: At first, low-quality cells and cell doublets were filtered out based on these criteria: 2% of cells with greater read-counts RNA, 2% of cells of each sample with low numbers of genes, or cells with more than 40% mitochondrial counts.

The filtered data were then normalized using ‘LogNormalize’ methods, and 2000 most variable genes were determined using the ‘FindVariableFeatures’ function in Seurat. Samples were then integrated using the ‘IntegrateData’ function from the Seurat package to correct the batch effect.

### Principal component analysis (PCA) and clustering

The TEs were temporarily removed from the count matrix to ensure that they do not affect our clustering and analyze them independently. After scaling data, PCA was performed using the RunPCA function in Seurat with default parameters.

The KNN graph was conducted based on the PCA-reduced data, and unsupervised clustering was performed using ‘FindNeighbors’ and ‘FindClusters,’ respectively. The TEs were added to clustered count matrix for further analysis.

### Differential expression (DE)

DE analysis was conducted on ‘RNA’ assays of count matrix, based on the ‘MAST’ (30) method using ‘FindAllMarkers,’ and cell type was determined using cell markers.

### Scoring pathways and correlation test

The gene sets of pathways that are presumably related to the immune system functions were retrieved from Gene Ontology (GO) (31, 32), wiki pathway (33), Kyoto Encyclopedia of Genes and Genomes (KEGG) (34-36), and Reactome (37) databases (Table S2). Average expression levels of these gene set in myeloid clusters and upregulated genes (logFC>0 and p.adjust <0.05) were calculated using ‘AddModuleScore’ in the Seurat package.

The correlation between these pathways and upregulated genes in neutrophil clusters was evaluated using a calculated score with the ‘corr. test’ function in R version 4.0.4.

Sc-RNA reads were aligned to ‘hg38 reference genome’ and ‘repeatmasker tracks’ using STAR and sc-TE aligners to count both genes and TEs. In the next step, count matrixes were filtered, batch effects were removed, and all matrixes were integrated together. Then, TEs were removed temporarily from the integrated matrix. After PCA and clustering, TEs were added, and DE analysis was conducted between all clusters. Finally, the correlation between DE TEs and some pathways was investigated.

## Results

To decipher the potential role of TE dysregulation in the immune deviation induced by the SARS-CoV-2 virus in COVID-19 patients, scRNA-seq data, including a total of 324135 cells in the BAL samples obtained from eight severe patients and four healthy donors, were reanalyzed (Table 1) (17, 22).

**Table 1.** GEO accession number, clinical status, and clinical outcome of samples reanalyzed.

| GEO Accession | Clinical outcome | Clinical status | Tissue |
| --- | --- | --- | --- |
| GSM4593888 | - | Healthy | BAL |
| GSM4593891 | - | Healthy | BAL |
| GSM4593890 | - | Healthy | BAL |
| GSM4593892 | - | Healthy | BAL |
| GSM4762143 | Alive | Severe COVID | BAL |
| GSM4762155 | Alive | Severe COVID | BAL |
| GSM4762159 | Alive | Severe COVID | BAL |
| GSM4762144 | Alive | Severe COVID | BAL |
| GSM4762140 | Dead | Severe COVID | BAL |
| GSM4762152 | Dead | Severe COVID | BAL |
| GSM4762150 | Dead | Severe COVID | BAL |
| GSM4762147 | Dead | Severe COVID | BAL |

Using KNN graph constructed on the PCA-reduced data, single cells were clustered and then labeled based on their canonical cell markers (Table S1). Average and percent expression of canonical cell markers were shown in all clusters of three populations (healthy, alive, and dead patients with severe COVID-19) (Fig. S1, S2, S3). Besides, to visualize these clusters, uniform manifold approximation and projection (UMAP) plots in these three populations were shown below.

This plot, which was obtained by UMAP visualization, demonstrates major cell types in BAL samples obtained from (a) healthy, (b) survived, and (c) dead patients. Each cell type is depicted by a specific color.

Based on the canonical markers, we identified major clusters including neutrophils, myeloid-derived suppressor cells (MDSCs), monocyte and macrophage, dendritic cells, lymphoid cells, and epithelial cells. Among all these clusters, MDSC revealed a distinct pattern for TE upregulation (Table S3-5). These clusters, differentially expressed a high number of TEs (logFC>0 and adj. *p*-value <0.05) (Fig. 3). Moreover, MDSC clusters had a higher frequency in infected patients.

### MDSCs reveal distinct immunological functions

MDSCs clusters were identified in survived and dead patients, as labeled in Fig. 2.

By scoring each pathway based on the expression levels of its genes, the correlation between the overexpressed genes in myeloid cell type clusters (LogFC>0 and adj. *p*..value<0.05) and the selected pathways was evaluated for the healthy, alive, and dead populations. The results of this analysis are presented in Table S6-8.

**Fig. 2.**
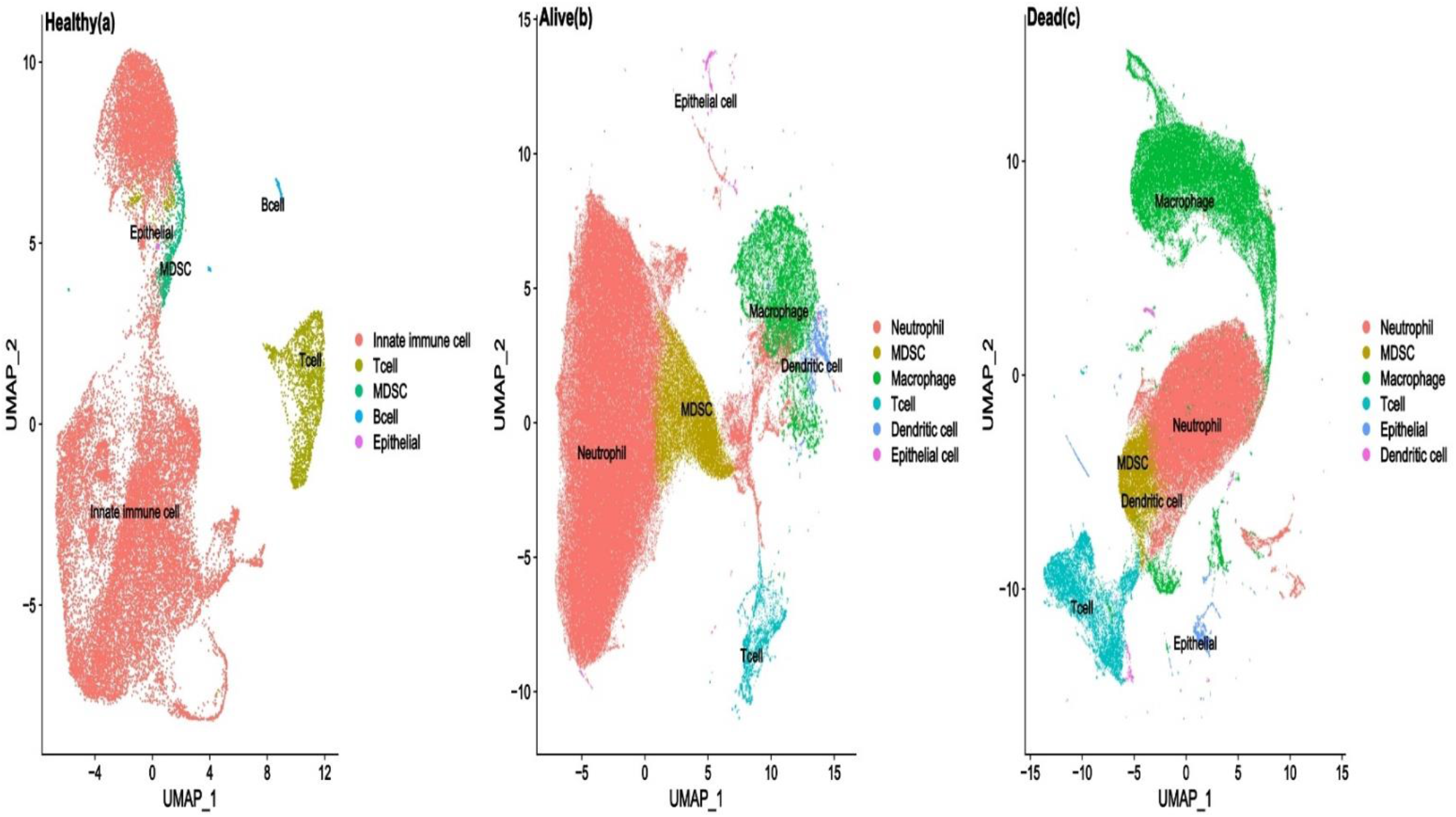
Single-cell analysis of cells from BAL samples of individuals with COVID-19 and controls.

**Fig. 3.**
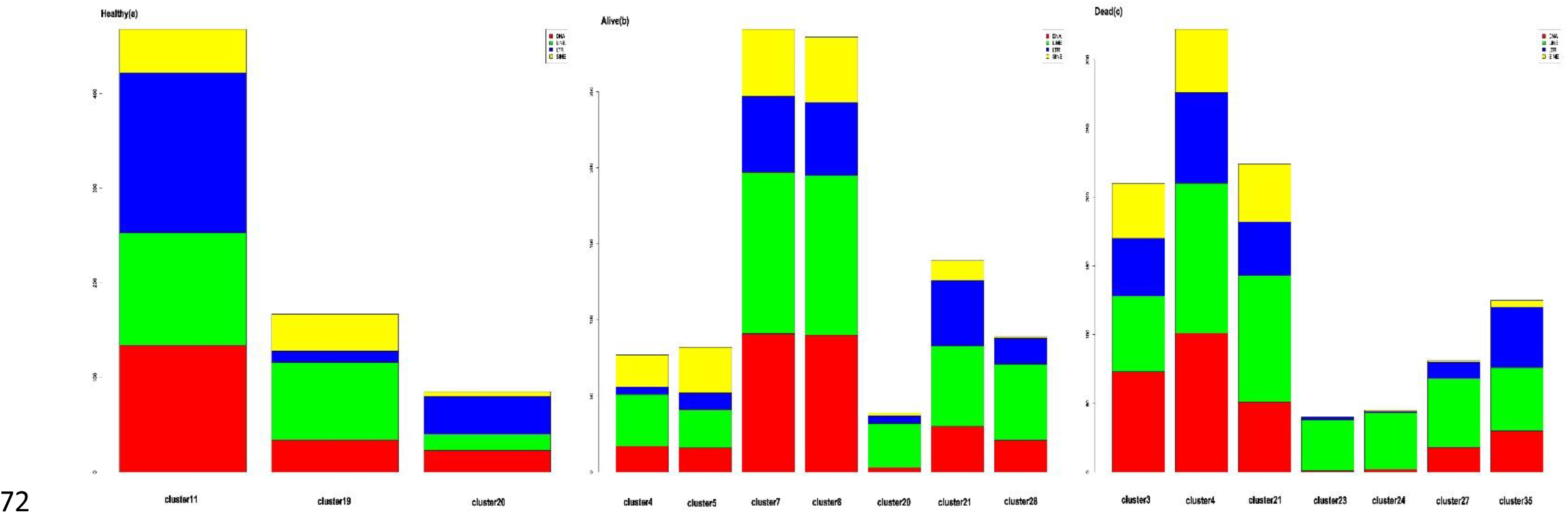
Bar plot summarizing the number of significantly upregulated TE subfamilies (logFC>0 and adj.*p*. value<0.05) in myeloid cell type clusters from healthy samples (a), alive (b), and dead (c) patients. The subfamilies of transposable elements include DNA transposons, long terminal repeat elements (LTR), long-interspersed elements (LINEs), and short-interspersed elements (SINEs) are depicted in red, blue, green, and yellow colors, respectively. The length of each bar represents the number of upregulated TEs from each subfamily. Clusters that have a TE number less than 50 are not represented in this illustration.

As demonstrated in Fig. S4-6 and Table S6-8, there is a significant difference in the function of MDSCs TEs and other myeloid cell type clusters. Some pathways related to neutrophil function were negatively associated with the function of MDSCs, which suggests the suppressive nature of these cells. On the other hand, some pathways including Toll receptor signaling pathway, TNF signaling pathway, and NF-kappa B signaling pathway were positively associated with the MDSCs function (Fig. 4).

**Fig. 4.**
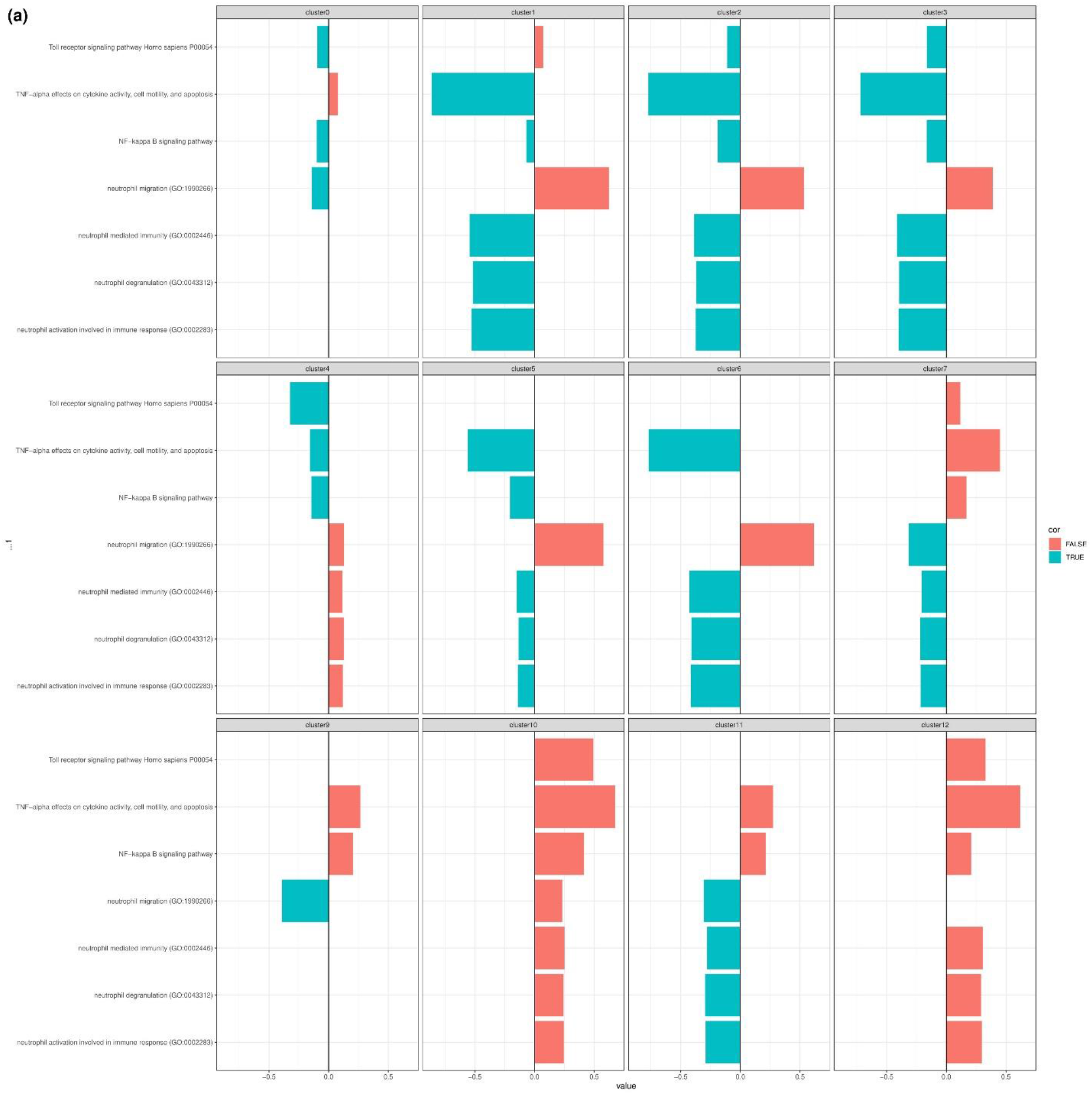
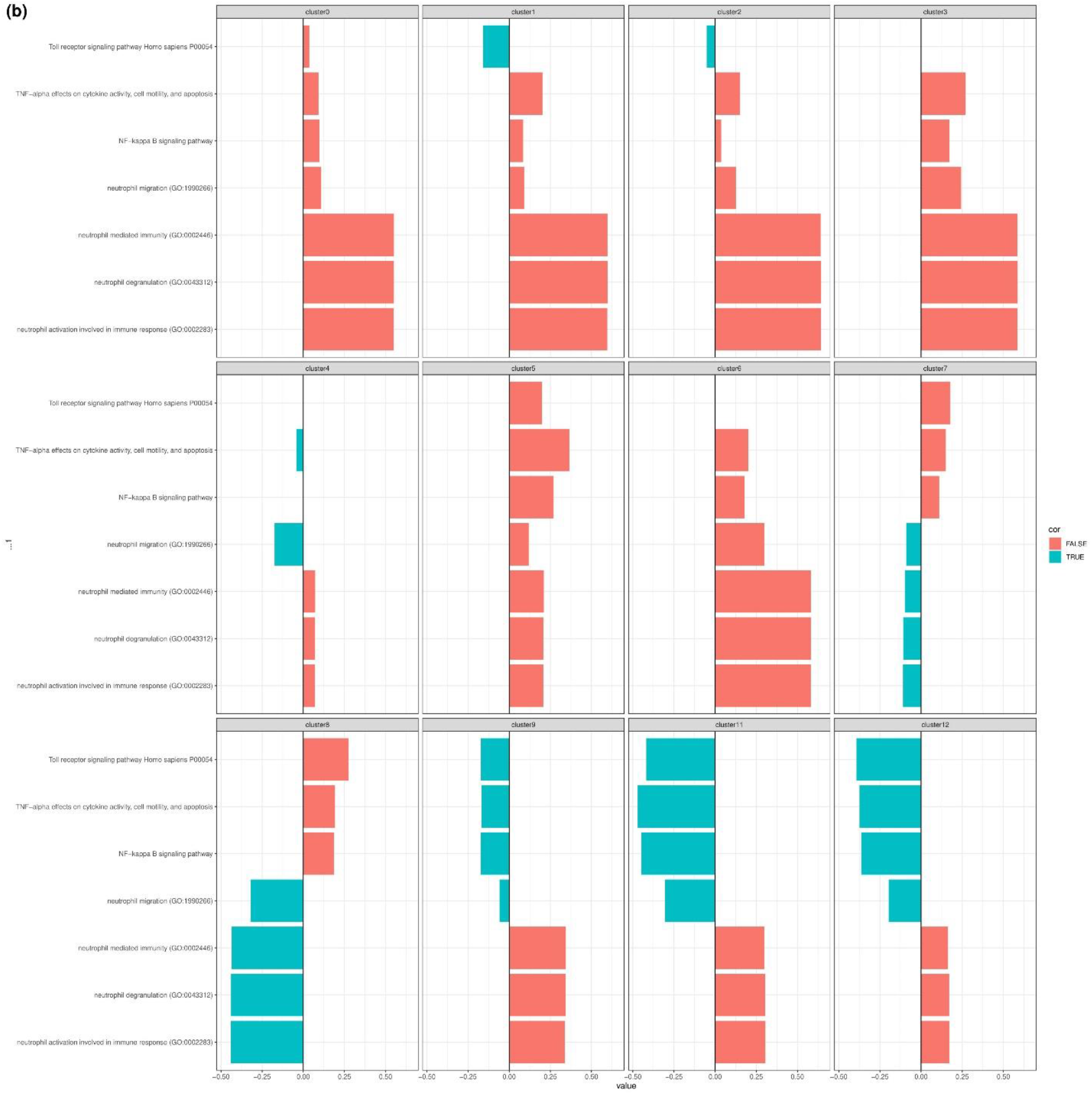
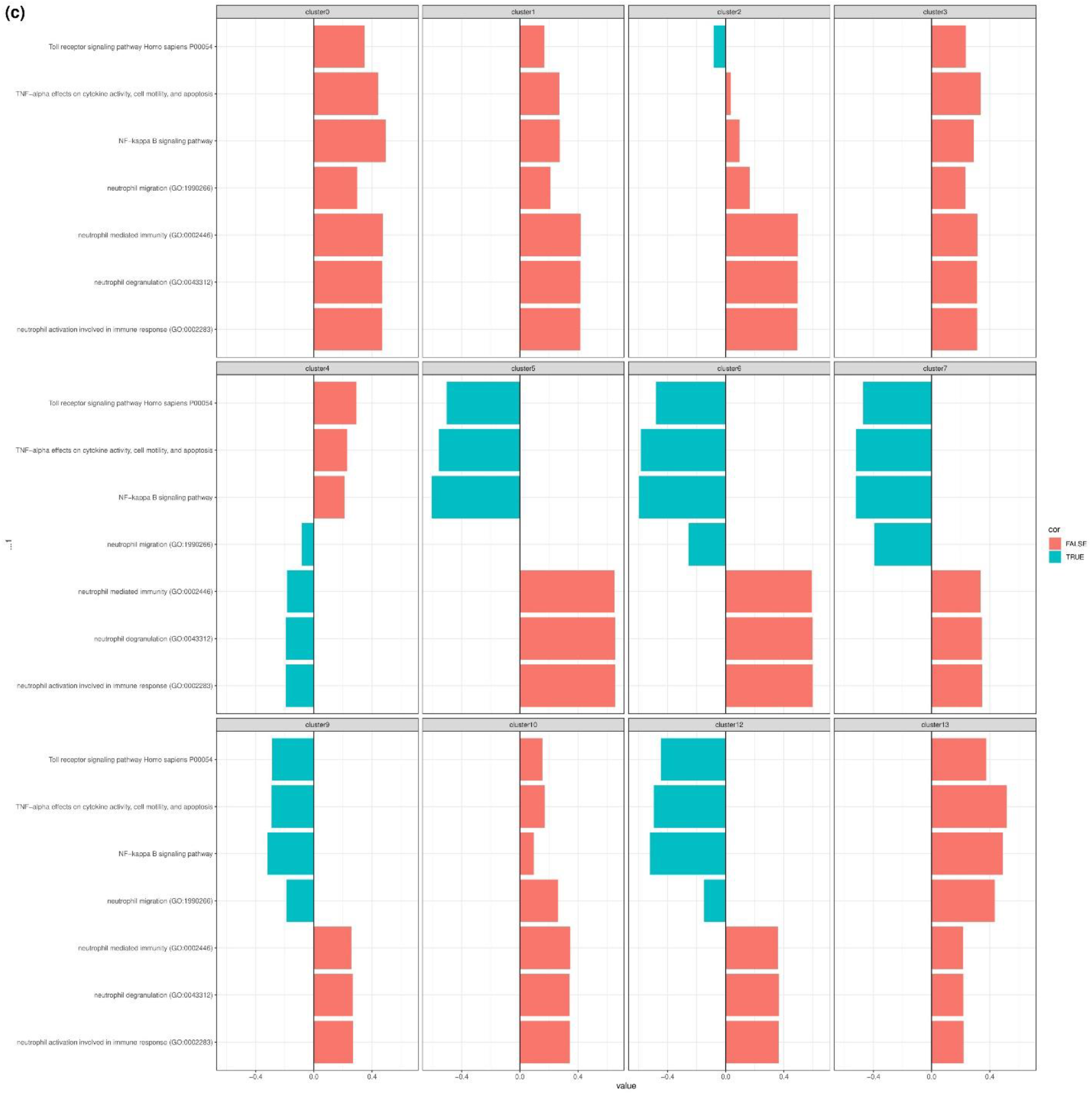
The correlation between five specific pathways and overexpressed genes in the first ten myeloid cell clusters from healthy, alive, and dead populations. There is a significant difference in the function of MDSC clusters with highly upregulated TEs compared to other myeloid cell types. The clusters with overexpressed TEs were found in cluster 11 of healthy samples (a), cluster 7 and 8 of alive patients (b), and cluster 4 of dead patients (c). MDSC clusters showed a negative correlation with neutrophil functions such as degranulation, activation involved in immune response, immunity, and migration. However, some pathways, including Toll receptor signaling pathway, TNF signaling pathway, and NF-kappa B signaling pathway, were positively associated with the function of MDSCs.

## Discussion

This study aimed to examine whether TE dysregulation correlates with the immune deviation induced by the SARS-CoV-2 virus in COVID-19 patients. To answer this question, the BAL sample obtained from 12 samples (including 4 control, 4 alive, and 4 dead patients) were reanalyzed at the single-cell level. We identified a cluster of MDSC with a higher population in COVID-19 than control.

Under normal homeostatic conditions, myeloid progenitors are produced in the bone marrow and appear as non-polarized or “resting” MDSCs with very low suppressive activity. These cells can migrate to the periphery and differentiate into mature macrophages, dendritic cells, and neutrophils, while losing their suppressive properties. However, under conditions of chronic inflammation, MDSCs including polymorphonuclear (PMN-MDSC) and monocytic (M-MDSC) ones undergo differentiation arrest, proliferation, and polarization towards highly suppressive cells that migrate to the periphery and sites of inflammation. Different subpopulations of MDSCs can sense changes in their surroundings and adjust their behavior accordingly. Therefore, in the context of chronic inflammation, the various subsets of MDSCs can sense changes in their environment and adapt accordingly due to their plastic nature. These cells can modify their developmental pattern, phenotype, and behavior in response to changes in environmental factors associated with chronic inflammation (38).

Research has shown that MDSCs play a critical role in regulating immune responses in various human diseases. MDSCs have the ability to inhibit T-cell proliferation and activation, modulate cytokine production by macrophages, suppress natural killer (NK) cell function, impair dendritic cell differentiation, and induce regulatory T cells (Tregs). Additionally, MDSCs can inhibit the proliferation and differentiation of B cells and induce regulatory B cells in multiple pathological conditions. These findings underscore the importance of MDSCs in the pathogenesis of diseases and highlight their potential as a therapeutic target (39-41).

Various studies have utilized advanced phenotypic and molecular techniques to investigate how the immune system interacts with the virus in COVID-19. Severe COVID-19 cases are characterized by alteration in the abundance, phenotype, and functionality of neutrophils. High numbers of neutrophils have been found in the nasopharyngeal epithelium, lungs, and blood of infected patients. Single-cell RNA sequencing has revealed the emergence of immature neutrophils that resemble PMN-MDSCs, suggesting the presence of immunosuppressive neutrophil precursors in severe COVID-19 cases. These precursors, may be released prematurely from the bone marrow and infiltrate the lung tissue in severe cases, leading to the expansion of PMN-MDSCs and contributing to the observed neutrophilia (42, 43).

We identified that TEs are highly upregulated in some clusters with the resemblance to myeloid cells. Notably, these cells were not recognized by canonical myeloid cell markers. These clusters were annotated as myeloid-derived suppressor cells based on the upregulation of some markers including Colony Stimulating Factor 3 Receptor(CSF3R) (44, 45), Nicotinamide Phosphoribosyltransferase(NAMPT) (46, 47), and nuclear paraspeckle assembly transcript 1 (NEAT1) (48, 49). The suppressive nature of MDSCs is in accordance with the obtained results suggestive of the negative association of canonical neutrophil functions including neutrophil migration, neutrophil-mediated immunity, neutrophil degranulation, and neutrophil activation involved in immune response with the MDSCs function (50).

Long non-coding RNAs (lncRNAs), including NEAT1 and metastasis-associated lung adenocarcinoma transcript 1 (MALAT1), were found to be highly upregulated in MDCS clusters with overexpressed TEs (Tables S8-S10). This result is confirmed by others who demonstrated that most lncRNAs are expressed under the control of TE promoters (51). Recent studies suggest that the upregulation of NEAT and MALAT1 may be associated with inflammation and consequent tissue damage seen in severe COVID-19 (52-54). Besides, it has been reported that TE overexpression is associated with inflammatory diseases. For instance, Macchietto et al. proved that TE overexpression is common in different viral infections (55). The overexpression of TEs in SARS-CoV-2 infection is confirmed as well (56). Noteworthy, autoantibodies against the endonuclease domain of the LINE1 gene are discovered in approximately 40% of SARS patients, which may be crucial in its pathogenesis (57).

Transcriptional regulatory networks are responsible for determining cellular identity, function, and response to stimuli by controlling gene expression programs. Regulatory elements such as promoters and enhancers act as “wires” in the genome, connecting genes into regulatory networks and regulating nearby gene expression (58). TEs have been suggested to play a role in the evolution of regulatory networks due to their ability to replicate throughout the host genome. While most TEs no longer encode functional proteins, they often retain transcription factor binding sites and can influence the expression of nearby genes. In recent years, there have been several studies demonstrating the important role of TEs in host gene regulation, leading to the conclusion that the cooption of TEs is a general mechanism for shaping the evolution of mammalian gene regulatory networks (59-61).

Recent research suggests that TEs are frequently coopted to regulate genes involved in immune processes. Studies have identified TEs being used as promoters (62), interferon-inducible enhancers (61), and insulator elements in immune cells (63). The unique pressures involved in the evolution of immune regulatory networks may favor the cooption of TEs. Immune genes are among the fastest evolving genes in the genome, reflecting the constant need to adapt to new and evolving pathogens (58, 61). TEs, which are a major source of genetic polymorphism, may facilitate rapid adaptive evolution of immune responses at the gene regulatory level due to their active or recently active status. In this regard, in a study by Ye et al. (4), chromatin profiling data from mouse CD8+ T lymphocytes was analyzed, revealing that multiple TE families contribute to predicted regulatory sequences. They also found that immune cells exhibit the highest enrichment of TE-derived enhancers compared to other cells, indicating that the cooption of TEs may more strongly influence immune regulatory networks. Thus, one may speculate that TE upregulation in MDSCs appears to be linked to the cells’ ability to adapt their phenotype and functional capabilities in response to changes in their microenvironment (38). TEs are repetitive and mobile elements, and their involvement in the genomic plasticity of MDSCs may occur through their insertion into coding or regulatory regions of immune-related genes. This can have a functional impact on gene expression, leading to plasticity (58, 64-66).

The defense against pathogens involves a series of coordinated events, beginning with the host’s recognition of the invading pathogen, in which Toll-like receptors (TLRs) play a crucial role. The purpose of sensing pathogens via TLRs is to quickly activate the innate immune response to eliminate the pathogen. Alveolar macrophages and neutrophils are well-known immune cells that are capable of phagocytosis and killing pathogens. Alveolar macrophages are usually the first responders, but they are later replaced by neutrophils, which are quickly recruited to the site of infection with the help of chemokines that are primarily produced by lung epithelial cells and macrophages. Neutrophils produce various harmful products, including reactive oxygen species and proteases, which can damage not only the pathogen but also the host’s own cells. After the pathogen has been eliminated, the host’s next priority is to initiate an appropriate anti-inflammatory response to prevent further neutrophil recruitment. Neutrophils have a short lifespan and begin to undergo apoptosis at the site of inflammation. To prevent lung injury, phagocytes must rapidly clear these apoptotic cells, a process known as efferocytosis. This is the stage at which MDSCs become involved in the host’s response to infection. MDSCs do not accumulate rapidly in the lung after infection. Instead, they develop late after infection, which makes sense; because lung MDSCs produce IL-10, which can impede neutrophil recruitment if produced too early. It has been revealed that lung MDSCs efficiently clear apoptotic neutrophils through efferocytosis, aided by the IL-10 produced by the MDSCs. Successful pathogen clearance, reduction in neutrophil infiltration, and elimination of dead neutrophils ultimately restore tissue homeostasis. As we revealed in Fig. 4, pathways related to neutrophil function were negatively associated with the function of MDSCs, which suggests these cells play an essential role in resolving lung inflammation by the inhibition of neutrophil functions (50).

The TLR2 and TLR4 signaling pathways induce the suppressive activity of MDSCs. Activation of the NF-κB pathway by TLR2/4 leads to the expression of inflammatory factors such as IL-6 and TNF-α. Subsequently, IL-6 and TNF-α activate both the STAT3 and NF-κB signaling pathways. Of particular note, the expression of the inflammatory factors S100A8 and S100A9 is regulated by STAT3. These factors act as TLR4 ligands, which then activate the NF-κB pathway, leading to an upregulation of IL-6 and TNF-α expression. This forms a feedback loop that enhances the expansion and activation of MDSCs (67).

As depicted in Fig. 4, some pathways including toll receptor signaling pathway, TNF signaling pathway, and NF-kappa B signaling pathway were positively associated with the MDSCs function. Since PRRs, the same as TLRs, could recognize TEs (68, 69), one may speculate that the upregulated TEs in MDSCs could enhance the suppressive activity of these cells. Although this assumption is based on limited evidence and requires further investigation to establish its validity (70), it serves as an initial proposition for further inquiry.

It should be noted that there were some limitations in our study. First of all, the number of samples that were reanalyzed was limited, and the lack of samples from mild patients in our analysis prevented us from generalizing this finding to COVID-19 mild patients. Besides, in this study, autonomous TE expression was not distinguished from co-transcription or pervasive transcription. This might lead to overestimating TE expression and its functional effects on the studied process (71).

All in all, this is the first study to examine the possible association of TE and its potential role in COVID-19 immune responses at the single-cell level. Our results suggest that the expansion of MDSCs could be a hallmark of COVID-19. Moreover, we recognized that TEs are highly upregulated in MDSCs. The upregulation of TEs in COVID-19 could be related to the plasticity of these cells in response to the microenvironments. Moreover, it seems that the recognition of overexpressed TEs by PRRs in MDSCs could strengthen the suppressive activity of these cells. Therefore, this study underscores the importance of TEs in the functionality of MDSCs in COVID-19. However, further studies are needed to decipher the probable causal link between TEs overexpression in MDSCs and their function seen in mild and severe COVID-19.

## Data Availability Statement

The datasets reanalyzed in this study were retrieved from the gene expression omnibus GEO database under accession numbers GSE157344 and GSE151928, respectively. Moreover, the code used in this study is available at git@github.com:mitra-frn/sc-TE-RNA.git.

## Author’s Contribution

Mitra Farahmandnejad, Pouria Mosaddeghi, Mohammadreza Dorvash, Pouya Faridi, and Manica Negahdaripour conceived the research idea. Mitra Farahmandnejad analyzed the data using the mentioned tools in the method section. Manica Negahdaripour, Amirhossein Sakhteman, and Pouya Faridi extensively reviewed the manuscript and analyzed the results. All authors wrote the manuscript and reviewed the last version of the manuscript.

## Acknowledgments

The authors wish to acknowledge CSC—IT Center for Science, Finland, for computational resources.

## References

1. Parasher A. COVID-19: Current understanding of its pathophysiology, clinical presentation and treatment. Postgraduate medical journal. 2021;97(1147):312–20.

2. Bagheri A, Moezzi SMI, Mosaddeghi P, Nadimi Parashkouhi S, Fazel Hoseini SM, Badakhshan F, et al. Interferon-inducer antivirals: Potential candidates to combat COVID-19. International Immunopharmacology. 2021;91:107245.

3. Parashkouhi SN, Mosaddeghi P, Bagheri A, Moezzi SMI, Hoseini SMF, Farahmandnejad M, et al. The dual sides of interferon induction in COVID-19 treatment. Trends in Pharmaceutical Sciences. 2021;7:9–14.

4. Wicker T, Sabot F, Hua-Van A, Bennetzen JL, Capy P, Chalhoub B, et al. A unified classification system for eukaryotic transposable elements. Nature Reviews Genetics. 2007;8(12):973–82.

5. Jönsson ME, Garza R, Johansson PA, Jakobsson J. Transposable Elements: A Common Feature of Neurodevelopmental and Neurodegenerative Disorders. Trends in Genetics. 2020;36(8):610–23.

6. Andrenacci D, Cavaliere V, Lattanzi G. The role of transposable elements activity in aging and their possible involvement in laminopathic diseases. Ageing research reviews. 2020;57:100995.

7. Larsen PA, Lutz MW, Hunnicutt KE, Mihovilovic M, Saunders AM, Yoder AD, et al. The Alu neurodegeneration hypothesis: A primate-specific mechanism for neuronal transcription noise, mitochondrial dysfunction, and manifestation of neurodegenerative disease. Alzheimer’s & Dementia. 2017;13(7):828–38.

8. Burns KH. Our conflict with transposable elements and its implications for human disease. Annual Review of Pathology: Mechanisms of Disease. 2020;15:51–70.

9. Zhang X, Zhang R, Yu J. New understanding of the relevant role of LINE-1 retrotransposition in human disease and immune modulation. Frontiers in cell and developmental biology. 2020:657.

10. Burns KH. Transposable elements in cancer. Nature Reviews Cancer. 2017;17(7):415–24.

11. Ferrarini MG, Lal A, Rebollo R, Gruber AJ, Guarracino A, Gonzalez IM, et al. Genome-wide bioinformatic analyses predict key host and viral factors in SARS-CoV-2 pathogenesis. Communications biology. 2021;4(1):1–15.

12. Kitsou K, Kotanidou A, Paraskevis D, Karamitros T, Katzourakis A, Tedder R, et al. Upregulation of Human Endogenous Retroviruses in Bronchoalveolar Lavage Fluid of COVID-19 Patients. Microbiology spectrum. 2021;9(2):e01260–21.

13. Mustafin R, Khusnutdinova E. COVID-19, Retroelements, and Aging. Advances in Gerontology. 2021;11(1):83–92.

14. El-Shehawi AM, Alotaibi SS, Elseehy MM. Genomic study of COVID-19 corona virus excludes its origin from recombination or characterized biological sources and suggests a role for HERVS in its wide range symptoms. Cytology and Genetics. 2020;54(6):588–604.

15. Zhang L, Richards A, Barrasa MI, Hughes SH, Young RA, Jaenisch R. Reverse-transcribed SARS-CoV-2 RNA can integrate into the genome of cultured human cells and can be expressed in patientderived tissues. Proceedings of the National Academy of Sciences. 2021;118(21):e2105968118.

16. Sorek M, Meshorer E, Schlesinger S. Transposable elements as sensors of SARS-CoV-2 infection. bioRxiv. 2021:2021.02.25.432821.

17. Bost P, De Sanctis F, Canè S, Ugel S, Donadello K, Castellucci M, et al. Deciphering the state of immune silence in fatal COVID-19 patients. Nature Communications. 2021;12(1):1428.

18. Stephenson E, Reynolds G, Botting RA, Calero-Nieto FJ, Morgan MD, Tuong ZK, et al. Single-cell multi-omics analysis of the immune response in COVID-19. Nature Medicine. 2021;27(5):904–16.

19. Xu G, Qi F, Li H, Yang Q, Wang H, Wang X, et al. The differential immune responses to COVID-19 in peripheral and lung revealed by single-cell RNA sequencing. Cell Discovery. 2020;6(1):73.

20. Edgar R, Domrachev M, Lash AE. Gene Expression Omnibus: NCBI gene expression and hybridization array data repository. Nucleic Acids Res. 2002;30(1):207–10.

21. Barrett T, Wilhite SE, Ledoux P, Evangelista C, Kim IF, Tomashevsky M, et al. NCBI GEO: archive for functional genomics data sets—update. Nucleic Acids Research. 2012;41(D1):D991–D5.

22. Mould KJ, Moore CM, McManus SA, McCubbrey AL, McClendon JD, Griesmer CL, et al. Airspace Macrophages and Monocytes Exist in Transcriptionally Distinct Subsets in Healthy Adults. American journal of respiratory and critical care medicine. 2021;203(8):946–56.

23. Kaminow B, Yunusov D, Dobin A. STARsolo: accurate, fast and versatile mapping/quantification of single-cell and single-nucleus RNA-seq data. bioRxiv. 2021:2021.05.05.442755.

24. He J, Babarinde IA, Sun L, Xu S, Chen R, Shi J, et al. Identifying transposable element expression dynamics and heterogeneity during development at the single-cell level with a processing pipeline scTE. Nature Communications. 2021;12(1):1456.

25. Fernandes JD, Zamudio-Hurtado A, Clawson H, Kent WJ, Haussler D, Salama SR, et al. The UCSC repeat browser allows discovery and visualization of evolutionary conflict across repeat families. Mobile DNA. 2020;11(1):13.

26. Hao Y, Hao S, Andersen-Nissen E, Mauck WM, III, Zheng S, Butler A, et al. Integrated analysis of multimodal single-cell data. Cell. 2021;184(13):3573–87.e29.

27. Stuart T, Butler A, Hoffman P, Hafemeister C, Papalexi E, Mauck WM, III, et al. Comprehensive Integration of Single-Cell Data. Cell. 2019;177(7):1888–902.e21.

28. Butler A, Hoffman P, Smibert P, Papalexi E, Satija R. Integrating single-cell transcriptomic data across different conditions, technologies, and species. Nature Biotechnology. 2018;36(5):411–20.

29. Satija R, Farrell JA, Gennert D, Schier AF, Regev A. Spatial reconstruction of single-cell gene expression data. Nature Biotechnology. 2015;33(5):495–502.

30. Finak G, McDavid A, Yajima M, Deng J, Gersuk V, Shalek AK, et al. MAST: a flexible statistical framework for assessing transcriptional changes and characterizing heterogeneity in single-cell RNA sequencing data. Genome Biology. 2015;16(1):278.

31. The Gene Ontology resource: enriching a GOld mine. Nucleic Acids Res. 2021;49(D1):D325–d34.

32. Ashburner M, Ball CA, Blake JA, Botstein D, Butler H, Cherry JM, et al. Gene ontology: tool for the unification of biology. The Gene Ontology Consortium. Nature genetics. 2000;25(1):25–9.

33. Martens M, Ammar A, Riutta A, Waagmeester A, Slenter DN, Hanspers K, et al. WikiPathways: connecting communities. Nucleic Acids Res. 2021;49(D1):D613–d21.

34. Kanehisa M. Toward understanding the origin and evolution of cellular organisms. Protein science : a publication of the Protein Society. 2019;28(11):1947–51.

35. Kanehisa M, Furumichi M, Sato Y, Ishiguro-Watanabe M, Tanabe M. KEGG: integrating viruses and cellular organisms. Nucleic Acids Res. 2021;49(D1):D545–d51.

36. Kanehisa M, Goto S. KEGG: kyoto encyclopedia of genes and genomes. Nucleic Acids Res. 2000;28(1):27–30.

37. Gillespie M, Jassal B, Stephan R, Milacic M, Rothfels K, Senff-Ribeiro A, et al. The reactome pathway knowledgebase 2022. Nucleic Acids Research. 2021;50(D1):D687–D92.

38. Ben-Meir K, Twaik N, Baniyash M. Plasticity and biological diversity of myeloid derived suppressor cells. Current Opinion in Immunology. 2018;51:154–61.

39. Lindau D, Gielen P, Kroesen M, Wesseling P, Adema GJ. The immunosuppressive tumour network: myeloid-derived suppressor cells, regulatory T cells and natural killer T cells. Immunology. 2013;138(2):105–15.

40. Li X, Zhong J, Deng X, Guo X, Lu Y, Lin J, et al. Targeting Myeloid-Derived Suppressor Cells to Enhance the Antitumor Efficacy of Immune Checkpoint Blockade Therapy. Front Immunol. 2021;12:754196.

41. Yang Y, Li C, Liu T, Dai X, Bazhin AV. Myeloid-Derived Suppressor Cells in Tumors: From Mechanisms to Antigen Specificity and Microenvironmental Regulation. Frontiers in Immunology. 2020;11.

42. Grassi G, Notari S, Gili S, Bordoni V, Casetti R, Cimini E, et al. Myeloid-Derived Suppressor Cells in COVID-19: The Paradox of Good. Frontiers in Immunology. 2022;13.

43. . !!! INVALID CITATION !!! {}.

44. Vetsika E-K, Koukos A, Kotsakis A. Myeloid-derived suppressor cells: major figures that shape the immunosuppressive and angiogenic network in cancer. Cells. 2019;8(12):1647.

45. Trikha P, Carson III WE. Signaling pathways involved in MDSC regulation. Biochimica et Biophysica Acta (BBA)-Reviews on Cancer. 2014;1846(1):55–65.

46. Wu Y, Pu C, Fu Y, Dong G, Huang M, Sheng C. NAMPT-targeting PROTAC promotes antitumor immunity via suppressing myeloid-derived suppressor cell expansion. Acta Pharmaceutica Sinica B. 2022;12(6):2859–68.

47. Travelli C, Consonni FM, Sangaletti S, Storto M, Morlacchi S, Grolla AA, et al. Nicotinamide Phosphoribosyltransferase Acts as a Metabolic Gate for Mobilization of Myeloid-Derived Suppressor CellsNAMPT Orchestrates Immunometabolism of MDSCs. Cancer research. 2019;79(8):1938–51.

48. Dong G, Yang Y, Li X, Yao X, Zhu Y, Zhang H, et al. Granulocytic myeloid-derived suppressor cells contribute to IFN-I signaling activation of B cells and disease progression through the lncRNA NEAT1-BAFF axis in systemic lupus erythematosus. Biochimica et Biophysica Acta (BBA) -Molecular Basis of Disease. 2020;1866(1):165554.

49. Liu Y, Han Y, Zhang Y, Lv T, Peng X, Huang J. LncRNAs has been identified as regulators of Myeloid-derived suppressor cells in lung cancer. Frontiers in Immunology. 2023;14.

50. Ray A, Chakraborty K, Ray P. Immunosuppressive MDSCs induced by TLR signaling during infection and role in resolution of inflammation. Frontiers in cellular and infection microbiology. 2013;3:52.

51. Lee H, Zhang Z, Krause HM. Long Noncoding RNAs and Repetitive Elements: Junk or Intimate Evolutionary Partners? Trends in genetics : TIG. 2019;35(12):892–902.

52. Huang K, Wang C, Vagts C, Raguveer V, Finn PW, Perkins DL. Long non-coding RNAs (lncRNAs) NEAT1 and MALAT1 are differentially expressed in severe COVID-19 patients: An integrated single-cell analysis. PloS one. 2022;17(1):e0261242.

53. Meydan C, Madrer N, Soreq H. The Neat Dance of COVID-19: NEAT1, DANCR, and Co-Modulated Cholinergic RNAs Link to Inflammation. Frontiers in Immunology. 2020;11.

54. Moazzam-Jazi M, Lanjanian H, Maleknia S, Hedayati M, Daneshpour MS. Interplay between SARS-CoV-2 and human long non-coding RNAs. J Cell Mol Med. 2021;25(12):5823–7.

55. Macchietto MG, Langlois RA, Shen SS. Virus-induced transposable element expression up-regulation in human and mouse host cells. Life science alliance. 2020;3(2).

56. Kitsou K, Kotanidou A, Paraskevis D, Karamitros T, Katzourakis A, Tedder R, et al. Upregulation of Human Endogenous Retroviruses in Bronchoalveolar Lavage Fluid of COVID-19 Patients. medRxiv. 2020.

57. He W-p, Shu C-l, Li B-a, Jun Z, Cheng Y. Human LINE1 endonuclease domain as a putative target of SARS-associated autoantibodies involved in the pathogenesis of severe acute respiratory syndrome. Chinese medical journal. 2008;121(7):608–14.

58. Ivancevic A, Chuong EB. Transposable elements teach T cells new tricks. Proceedings of the National Academy of Sciences. 2020;117(17):9145–7.

59. Fuentes DR, Swigut T, Wysocka J. Systematic perturbation of retroviral LTRs reveals widespread long-range effects on human gene regulation. eLife. 2018;7:e35989.

60. Römer C, Singh M, Hurst LD, Izsvák Z. How to tame an endogenous retrovirus: HERVH and the evolution of human pluripotency. Current opinion in virology. 2017;25:49–58.

61. Chuong EB, Elde NC, Feschotte C. Regulatory evolution of innate immunity through co-option of endogenous retroviruses. Science (New York, NY). 2016;351(6277):1083–7.

62. Romanish MT, Lock WM, de Lagemaat LNv, Dunn CA, Mager DL. Repeated recruitment of LTR retrotransposons as promoters by the anti-apoptotic locus NAIP during mammalian evolution. PLoS genetics. 2007;3(1):e10.

63. Wang J, Vicente-García C, Seruggia D, Moltó E, Fernandez-Miñán A, Neto A, et al. MIR retrotransposon sequences provide insulators to the human genome. Proceedings of the National Academy of Sciences. 2015;112(32):E4428–E37.

64. Chénais B. Transposable Elements and Human Diseases: Mechanisms and Implication in the Response to Environmental Pollutants. International journal of molecular sciences. 2022;23(5).

65. Bhat A, Ghatage T, Bhan S, Lahane GP, Dhar A, Kumar R, et al. Role of Transposable Elements in Genome Stability: Implications for Health and Disease. International Journal of Molecular Sciences. 2022;23(14):7802.

66. Mosaddeghi P, Farahmandnejad M, Zarshenas MM. The role of transposable elements in aging and cancer. Biogerontology. 2023.

67. Zhou H, Jiang M, Yuan H, Ni W, Tai G. Dual roles of myeloid-derived suppressor cells induced by Toll-like receptor signaling in cancer. Oncology letters. 2021;21(2):149.

68. Gazquez-Gutierrez A, Witteveldt J S RH, Macias S. Sensing of transposable elements by the antiviral innate immune system. RNA (New York, NY). 2021;27(7):735–52.

69. O’Donnell T, Vabret N. Repeat elements amplify TLR signaling. Nature Reviews Immunology. 2021;21(12):760–.

70. Clapes T, Polyzou A, Prater P, Sagar Morales-Hernández A, Ferrarini MG, et al. Chemotherapyinduced transposable elements activate MDA5 to enhance haematopoietic regeneration. Nature Cell Biology. 2021;23(7):704–17.

71. Lanciano S, Cristofari G. Measuring and interpreting transposable element expression. Nature reviews Genetics. 2020;21(12):721–36.

