## Supplemental Table 1 & 2 for "Correlation of myeloid-derived suppressor cell expansion with upregulated transposable elements in severe COVID-19 unveiled in single-cell RNA sequencing reanalysis": supp2-new.docx

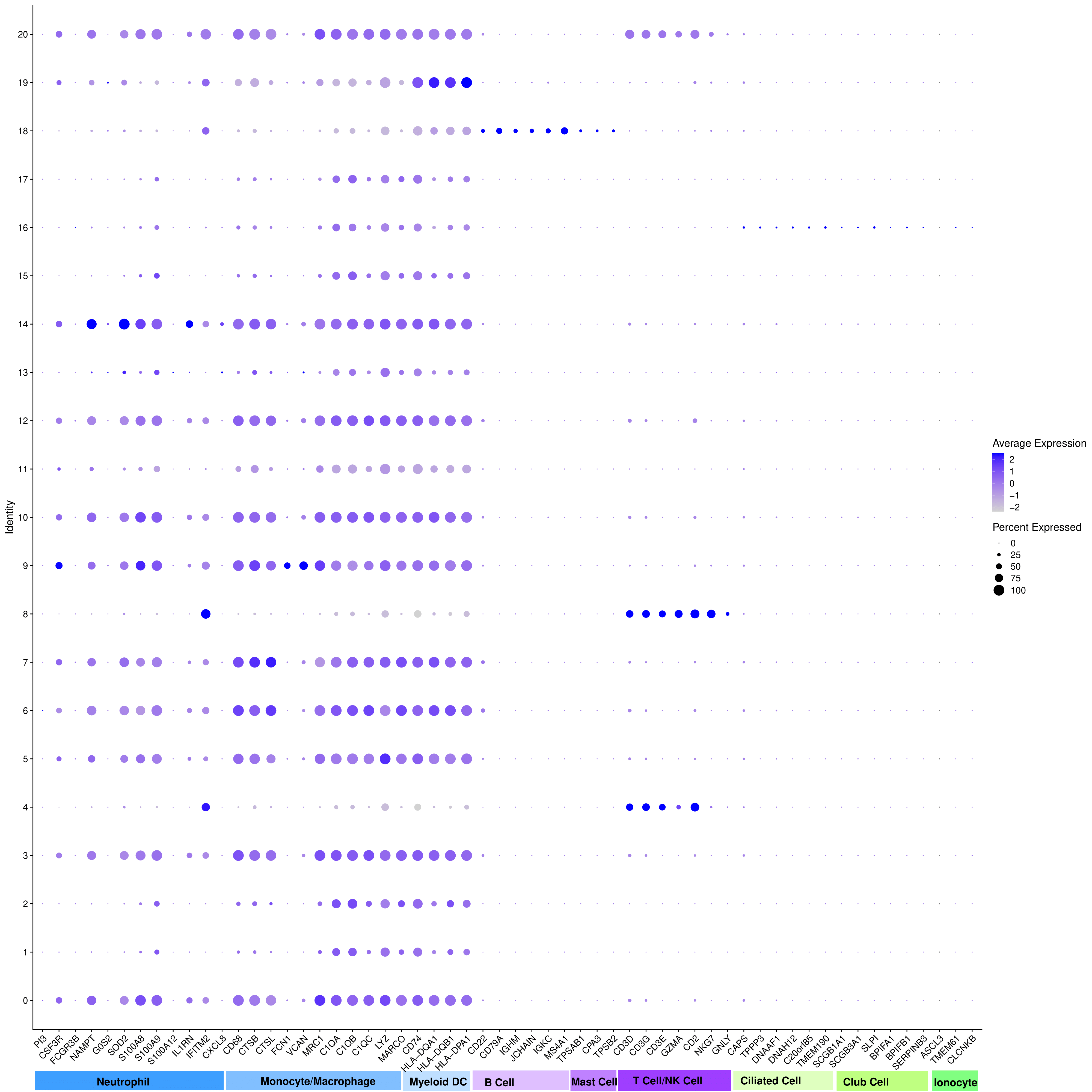

**Fig. S1**. Dot plot of canonical marker gene expression in the healthy population.

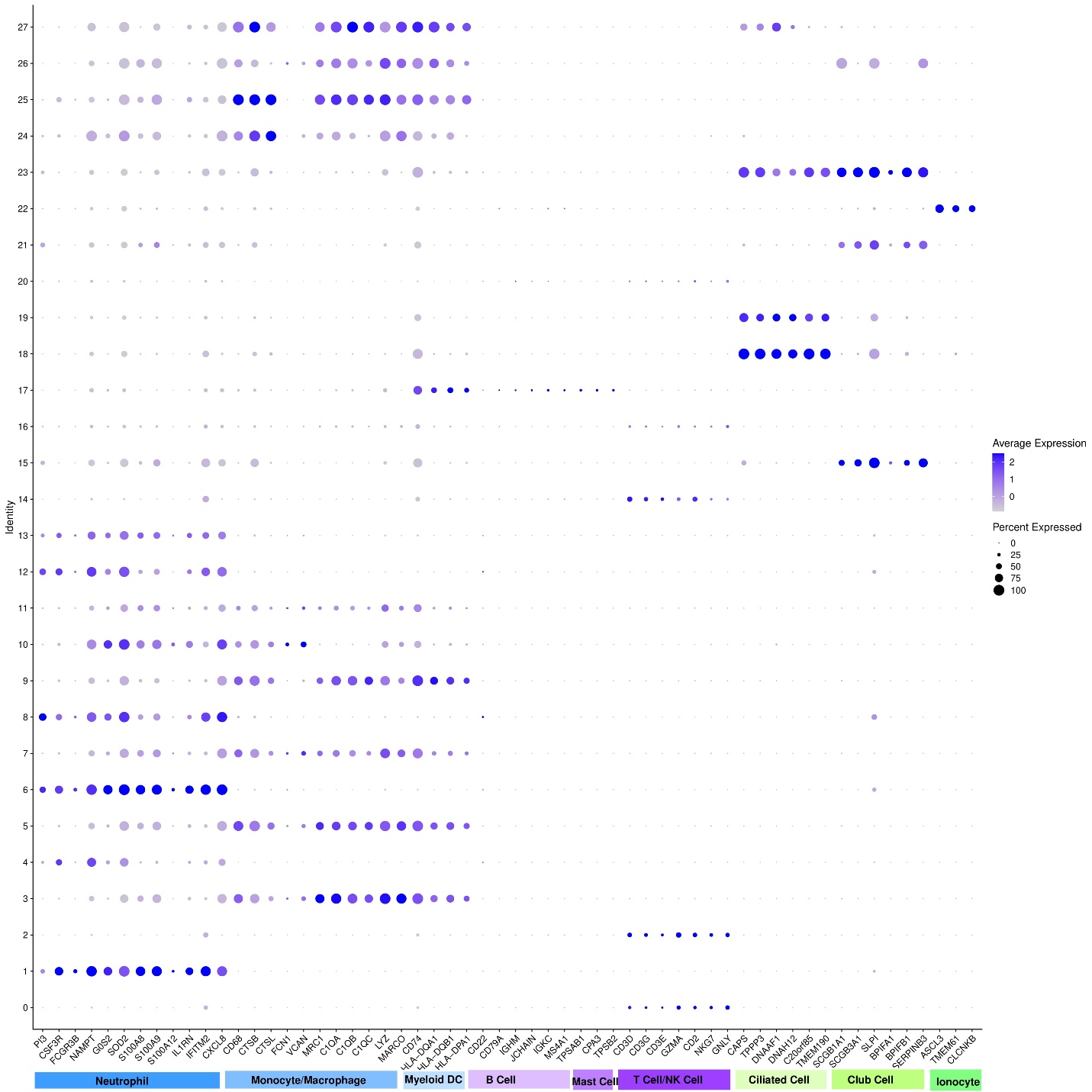
 **Fig. S2**. Dot plot of canonical marker gene expression in alive population.

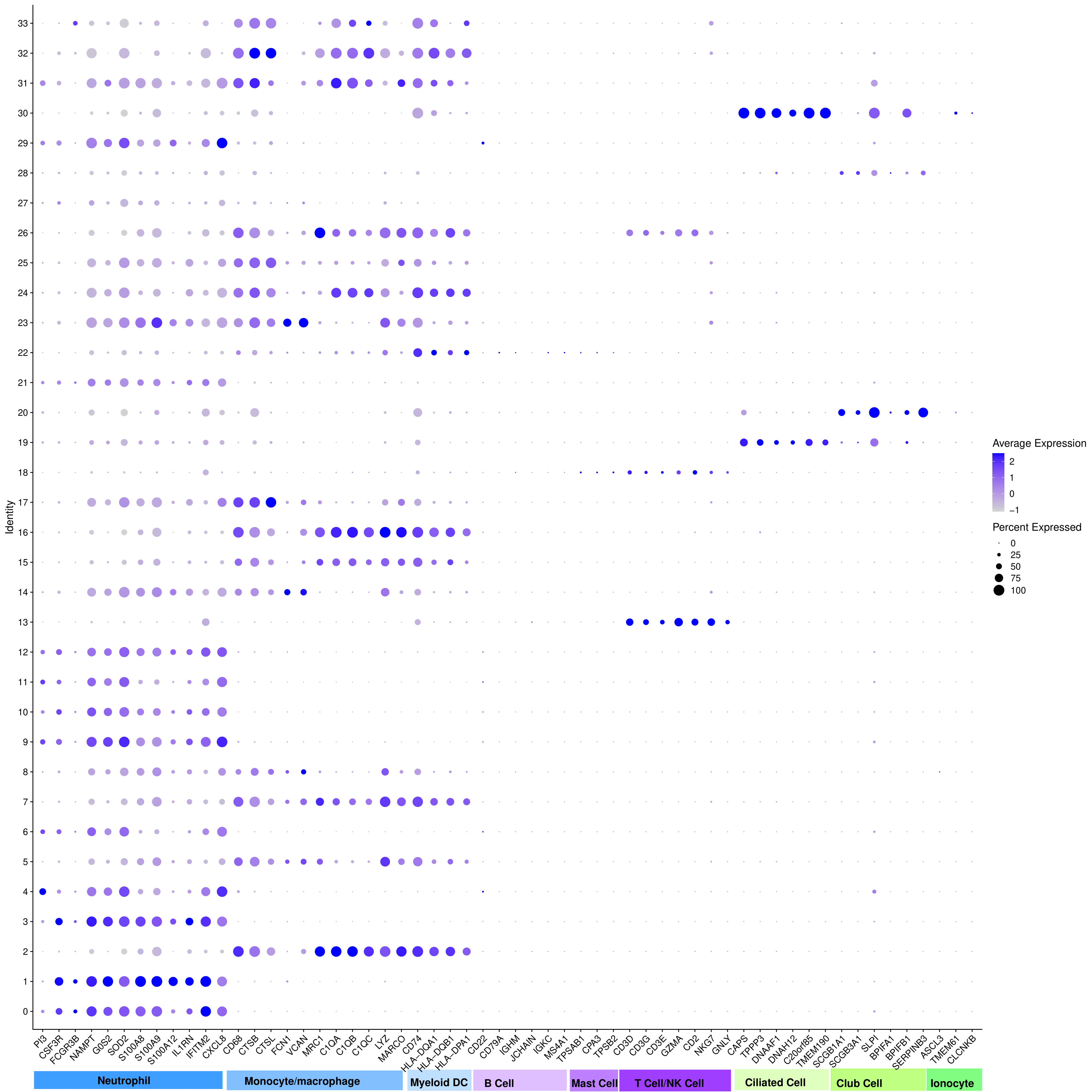

**Fig. S3**. Dot plot of canonical marker gene expression in dead population.

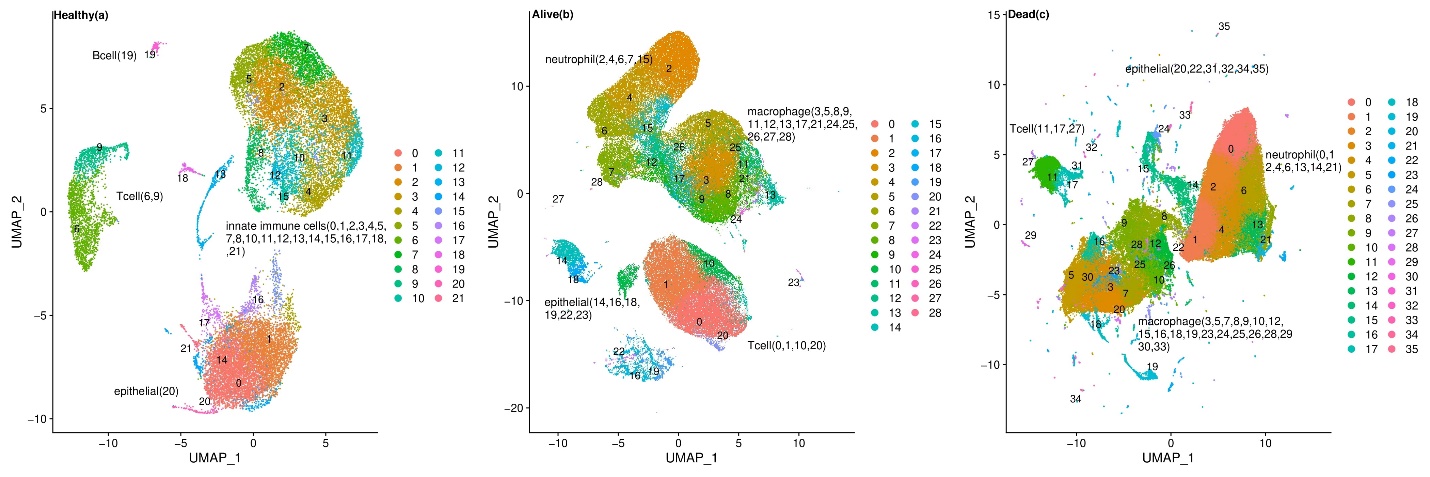
**Fig. S4.** UMAP visualization of primary cell types and associated clusters in BAL samples obtained from (a) healthy, (b) survived, and (c) dead patients in nointron mode.

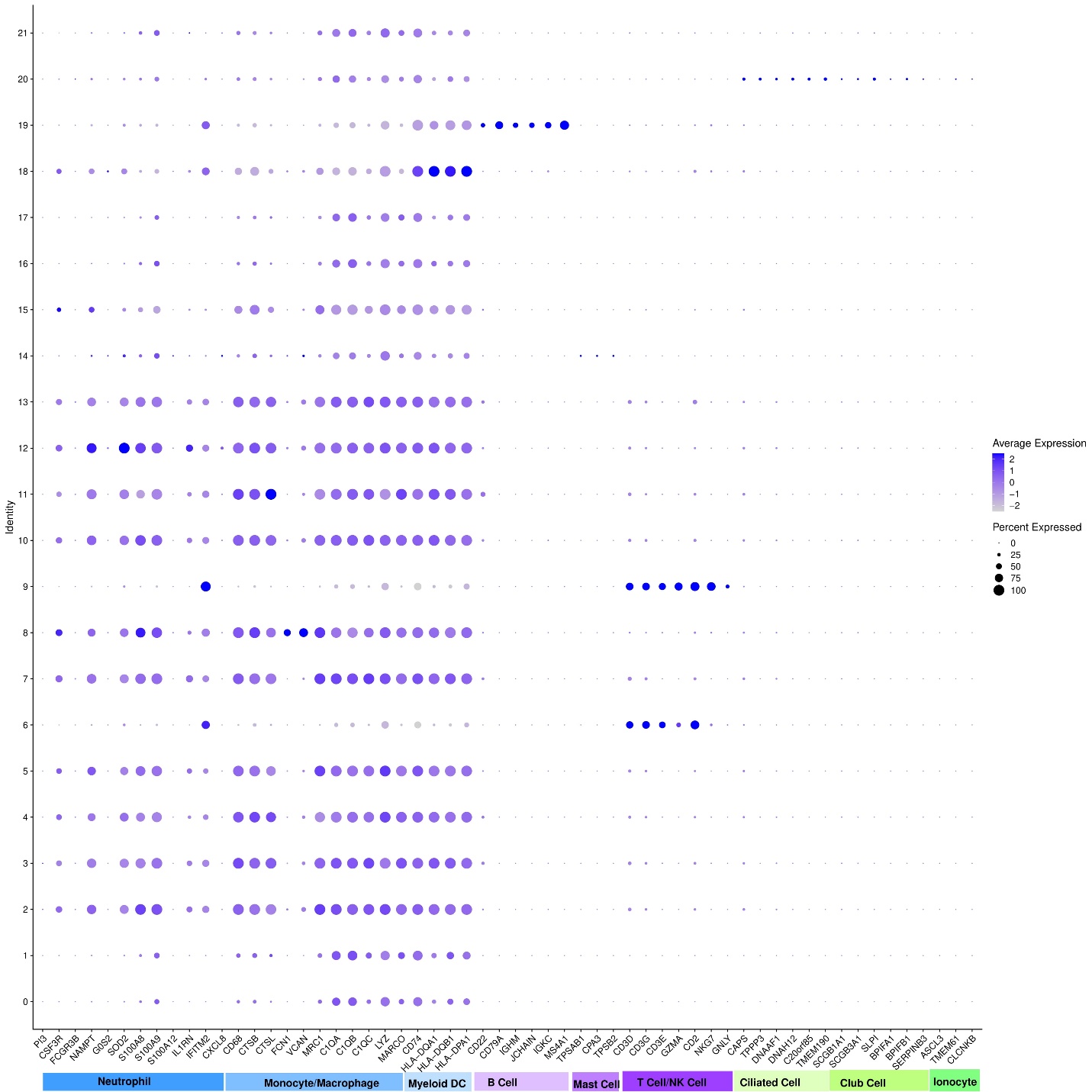

**Fig. S5**. Dot plot of canonical marker gene expression in healthy population in nointron mode

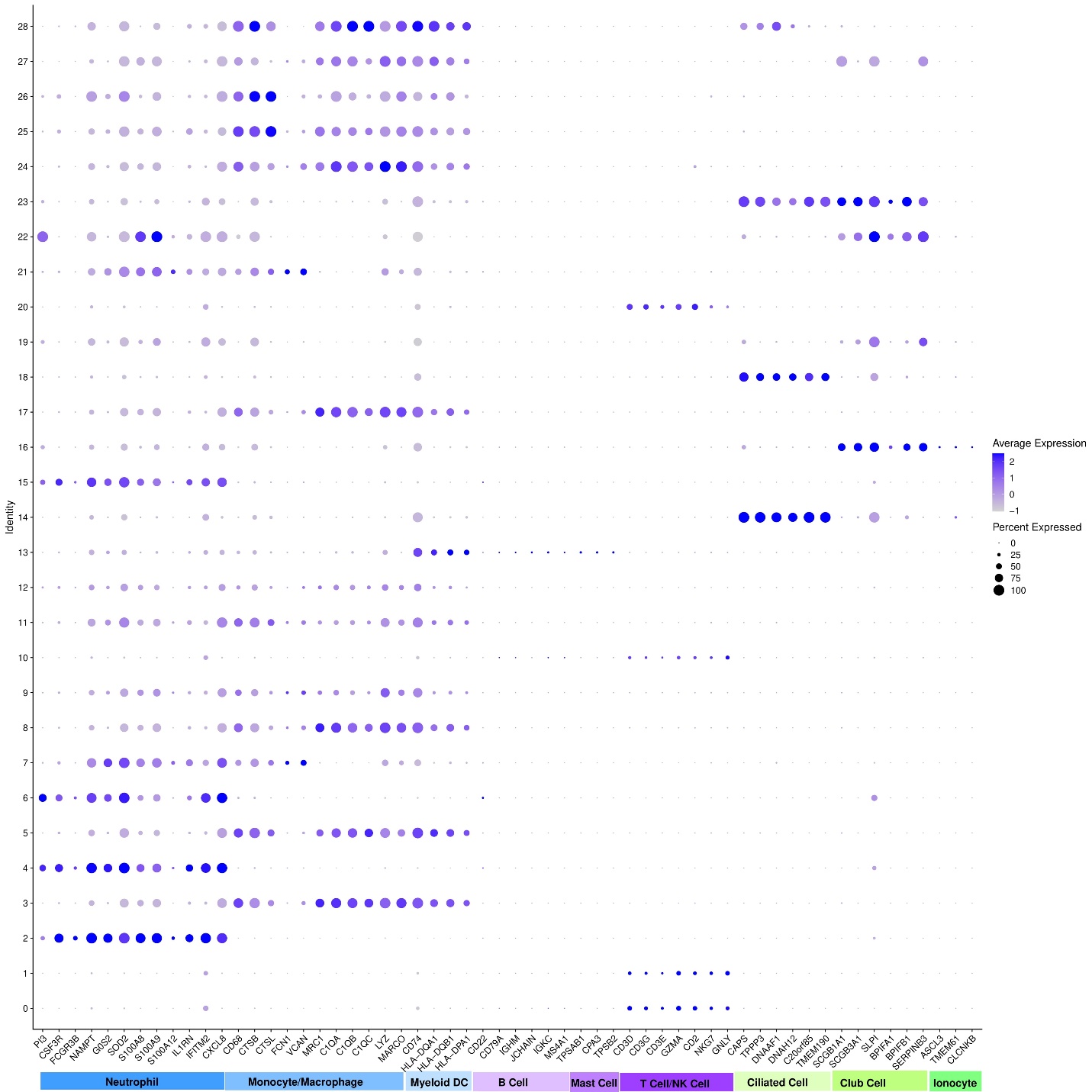

**Fig. S6**. Dot plot of canonical marker gene expression in alive population in nointron mode

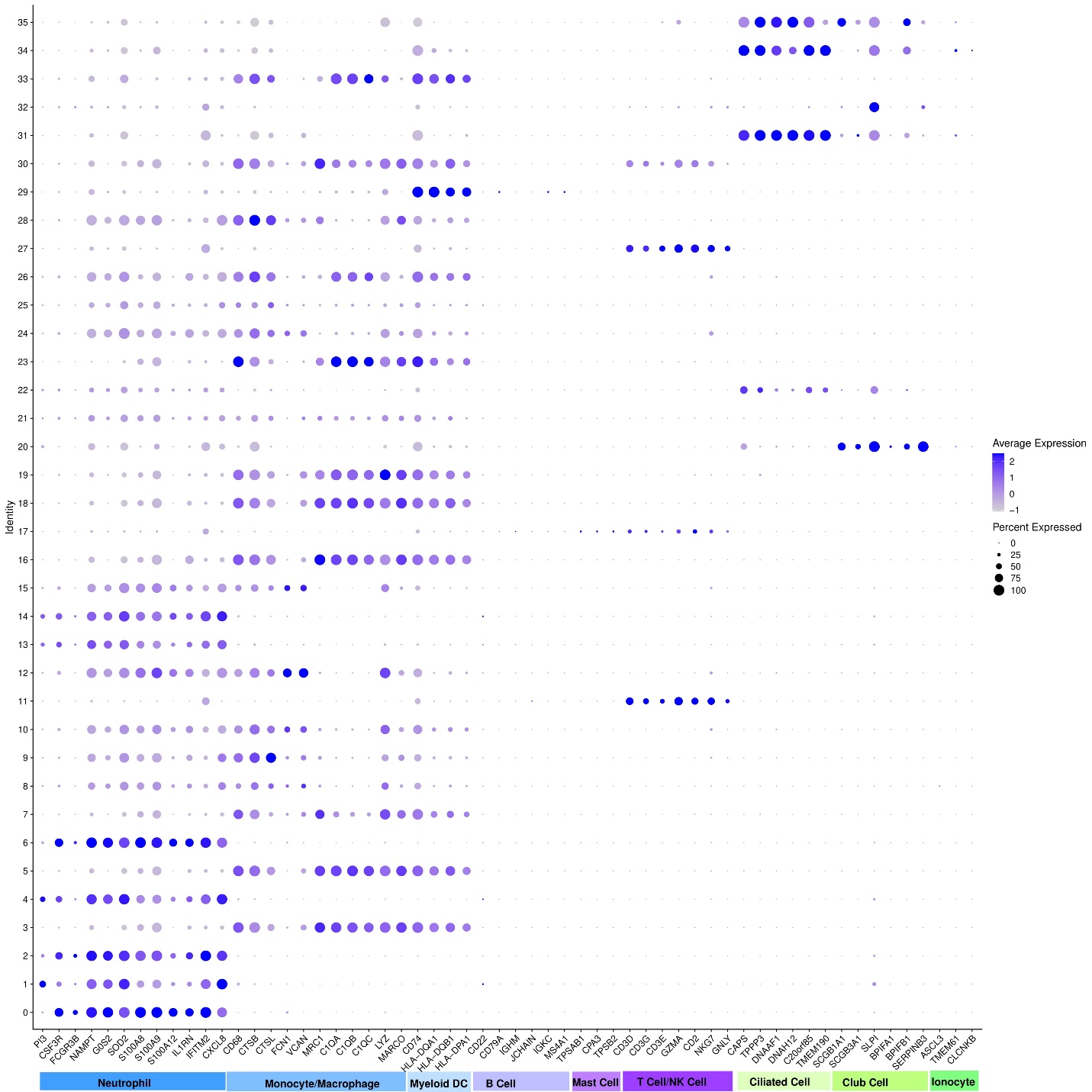

**Fig. S7**. Dot plot of canonical marker gene expression in dead population in nointron mode

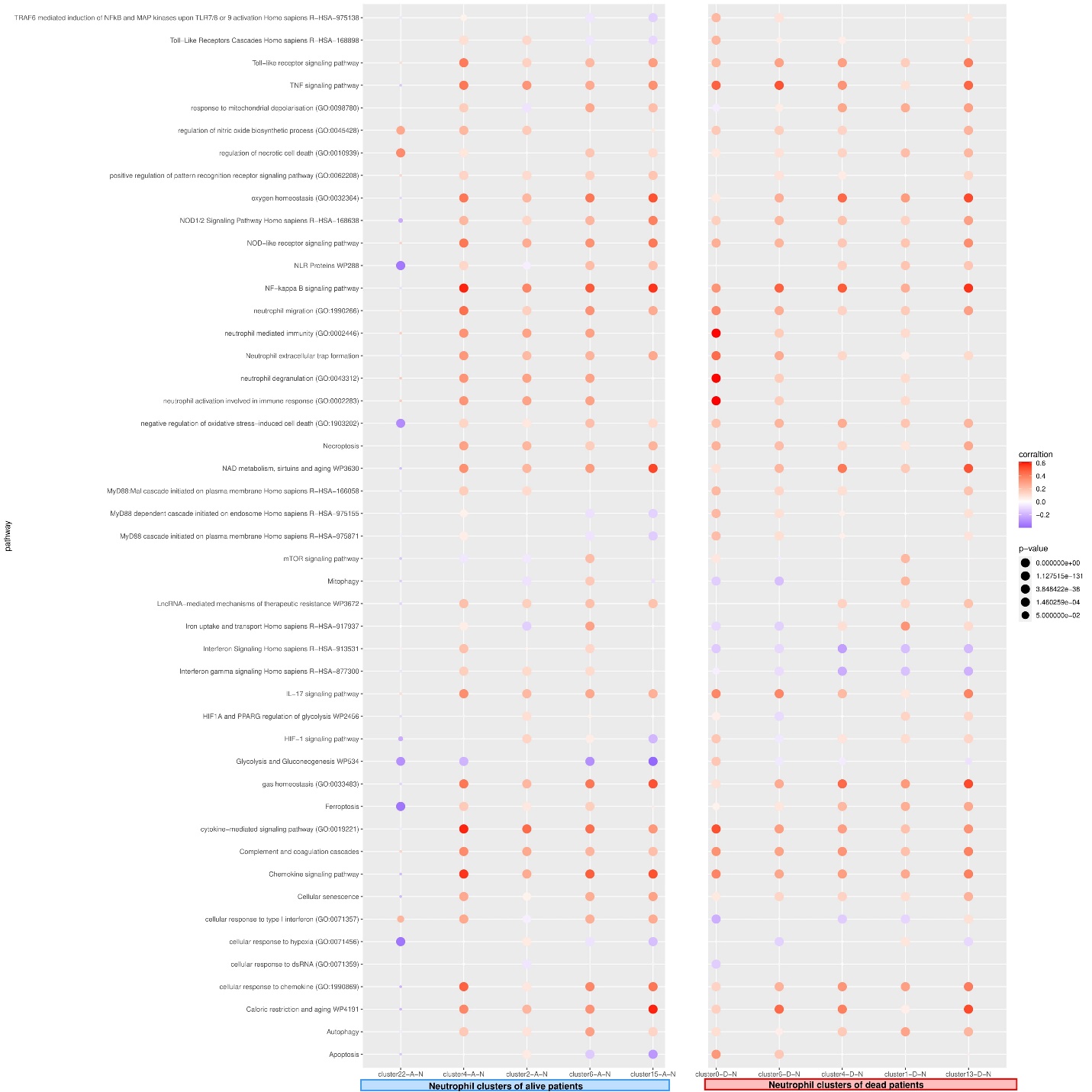

**Fig. S8.** Dot plot representing the probable relation of 46 selected gene sets with neutrophil function.
